# Enhanced late blight resistance by engineering an EpiC2B-insensitive immune protease

**DOI:** 10.1101/2023.05.29.541874

**Authors:** Mariana Schuster, Sophie Eisele, Liz Armas-Egas, Till Kessenbrock, Jiorgos Kourelis, Markus Kaiser, Renier A. L. van der Hoorn

## Abstract

Crop protection strategies relying on the improvement of the natural plant immune system via genetic engineering are sustainable solutions against the pathogen thread on food security. Here we describe a novel way to improve the plant immune system by immune protease engineering. As proof of concept, we increased resistance against the late blight pathogen *Phytopththora infestans* by rendering the tomato secreted immune protease Pip1 insensitive to the *P. infestans*-secreted inhibitor Epic2B. This concept can be applied to secreted immune proteases in crops by precision breeding.

---

Papain-like immune proteases (PLCPs) have emerged as promising engineering targets for crop protection, given their significant roles in plant immunity for key crops such as tomato, maize, and citrus (Misas-Villamil et al., 2016). The fact that they are targeted by a wide range of pathogen-secreted inhibitors highlights the importance of these proteases in defending against various pathogens. Depletion of the apoplastic immune PLCP *Phytopthora*-inhibited protease 1 (Pip1) from tomato, for instance, causes hyper-susceptibility to bacterial, fungal and oomycete tomato pathogens (Ilyas et al., 2015). Immunity by Pip1 in wild-type tomato is, however, suboptimal since Pip1 is suppressed during infection by diverse pathogen-secreted inhibitors, such as the cystatin-like EpiC2B from the oomycete late blight pathogen *Phytophthora infestans* (Tian et al., 2007). Here, we tested if we could increase Pip1-based immunity against late blight by engineering Pip1 into an EpiC2B-insensitive protease. To guide Pip1 mutagenesis, we generated a structural model of the EpiC2B-Pip1 complex using AlphaFold-Multimer (Evans et al., 2022). These structural models represent a classic interaction between the tripartite wedge of cystatin (EpiC2B) in the substrate binding groove of papain (Pip1). This model indicated that engineering Pip1 to prevent inhibition without affecting Pip1 substrate specificity is possible because the interaction surface of Pip1 with EpiC2B is larger than the substrate binding groove (**Figure 1A**).

**Figure 1.**
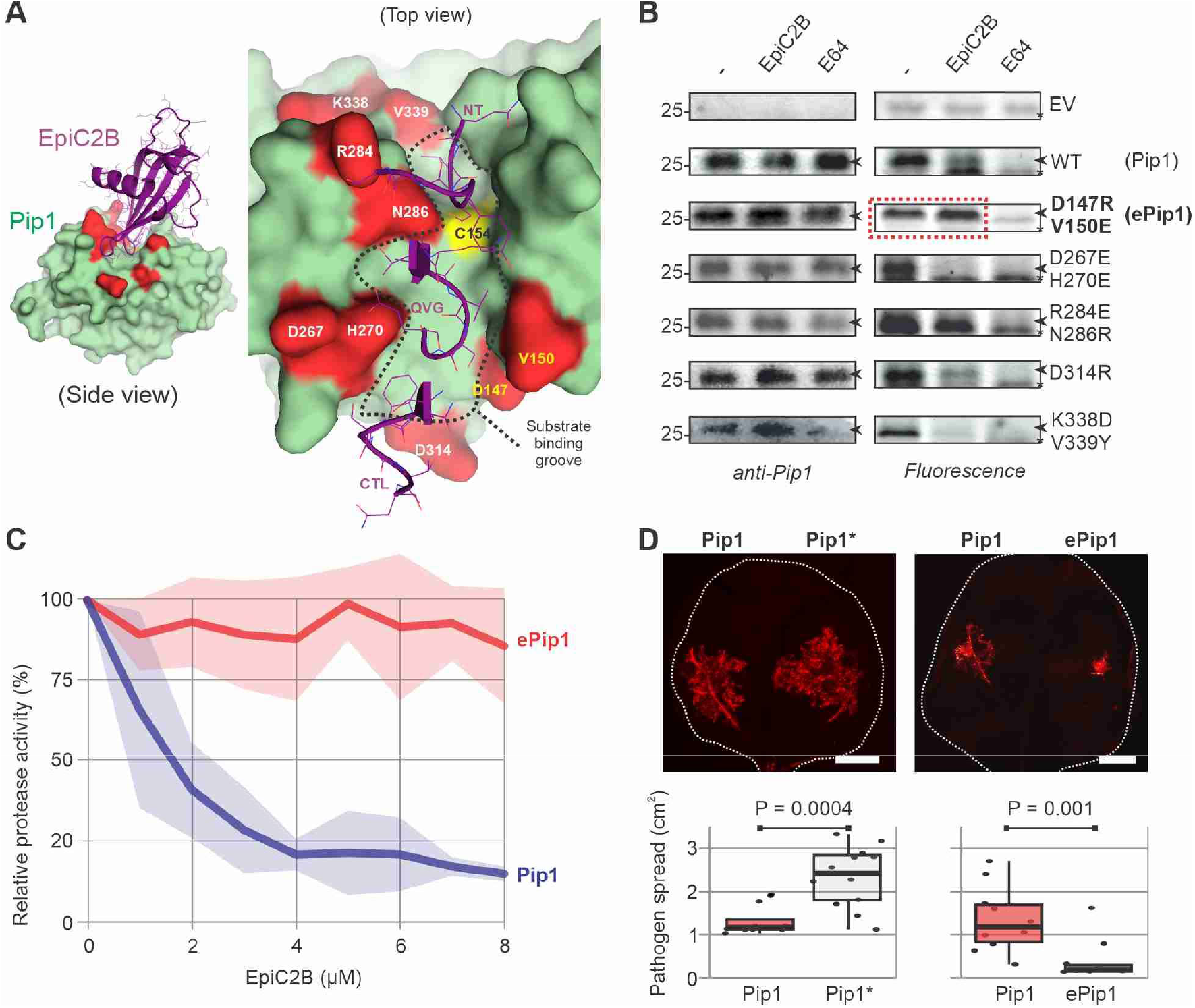
Engineered, EpiC2B-insensitive Pip1 reduces growth of *P. infestans* on *N. benthamiana*. **(A)** Structural model of the Pip1-EpiC2B complex generated with AlphaFold Multimer (Evans et al., 2022) showing the substrate binding groove (dashed line), and the residues selected for mutagenesis in Pip1 (red). The right zoomed image shows the tripartite loops of EpiC2B interacting with the substrate binding groove: the N-terminus (NT), the middle loop containing the QxVxG (QVG) motif and the C-terminal loop (CTL) containing a conserved tryptophane. **(B)** Pip1 mutants are active proteases with altered sensitivities to EpiC2B inhibition. Apoplastic fluids isolated from agroinfiltrated leaves transiently expressing (mutant) Pip1 were pre-incubated for 30 min with and without 100 µM E-64 or 3 µM EpiC2B, and then labelled for 3 hours with 0.2 µM TK011. Samples were separated on SDS-PAGE gels, scanned for fluorescence (right panel) and then transferred onto a membrane for an α-Pip1 western blot (left panel). **(C)** Two amino acid substitutions render Pip1 insensitive to EpiC2B. Apoplastic fluids isolated from agroinfiltrated leaves transiently expressing Pip1 and engineered ePip1 were pre-incubated for 30 min with and without 100 µM E-64, or increasing concentrations of EpiC2B, and then labelled for 3 hours with 0.2 µM TK011. Samples were separated on SDS-PAGE gels and scanned for fluorescence. Signal intensity was quantified using the Fiji platform (Schindelin et al., 2012). Values are depicted as a percentage of the labelling detected without inhibitor. Data are expressed as mean ± SE of n=3 replicates. **(D)** Engineered Pip1 restricts *P. infestans* growth in *N. benthamiana. N. benthamiana* leaf halves where agroinfiltrated with wild-type Pip1, catalytically dead Pip1* or engineered ePip1 for pairwise comparisons. Three days after agroinfiltration, the leaves where detached and zoospores of *P. infestans* strain 88069 tD was drop-inoculated onto each leaf half. Leaves where incubated for further 7 to 9 days and imaged for fluorescence. The area containing fluorescent hyphae was measured using the Fiji platform. P values correspond to a paired *t*-test of n>8 replicates. Results shown correspond to a representative example of 3 biological replicates (see Fig S1).

We selected nine residues for targeted mutagenesis of Pip1 that are predicted to directly interact with EpiC2B but are not in the substrate binding groove (**Figure 1A**). To generate the greatest disruption in protease-inhibitor interaction, we substituted these residues into bulky amino acids of the opposite charge. Following this strategy, we generated four double mutants and one single mutant of Pip1 and produced the proteins *in planta* via agroinfiltration. All proteins were detected in the apoplast of agroinfiltrated *Nicotiana benthamiana* plants using the anti-Pip1 antibody (**Figure 1B**). All Pip1 proteases are active proteases because they reacted with TK011, which is a fluorescent activity-based probe for PLCPs (Supplemental **Figure S1**), generated by coupling azide-E64 with alkyne-Bodipy using click chemistry. Preincubation with 3 µM purified EpiC2B reduced labelling of Pip1 and some of the Pip1 mutants. Importantly, the D147R/V150E mutant of Pip1 seems insensitive to EpiC2B inhibition (**Figure 1B**) and was hence called engineered Pip1 (ePip1). Preincubation with increasing EpiC2B concentrations revealed that ePip1 is insensitive to up to 8 μM EpiC2B, whereas wild-type Pip1 can be inhibited by 1 μM EpiC2B (**Figure 1C**).

We next tested if ePip1 enhances immunity against late blight. Taking advantage of the fact that *N. benthamiana* is a natural null mutant for Pip1 (Kourelis et al., 2020), and that infections of agroinfiltrated leaves are well-established assays for *P. infestans*, we first tested if transient expression of wild-type Pip1 enhances late blight immunity. A catalytic mutant of Pip1 (Pip1*), which lacks the two Cys residues in the active site (C153A/C154A), was included as a negative control. Wild-type Pip1 and mutant Pip1* were transiently expressed side-by-side in *N. benthamiana* leaves, and the agroinfiltrated area was infected with zoospores of transgenic *P. infestans* expressing fluorescent tDtomato (Chaparro-Garcia et al., 2011). The spread of the fluorescent pathogen was measured once the hyphae had grown beyond the inoculation droplet but within the agroinfiltrated zone (between 8 and 11 dpi) and quantified using the Fiji platform (Schindelin et al., 2012). The reduced fluorescent hyphae in tissue expressing Pip1 showed that wild-type Pip1 increases immunity against *P. infestans* growth when compared to the inactive mutant Pip1* negative control (**Figure 1D**), confirming that Pip1 is an immune protease, also when expressed in *N. benthamiana*. Having a transient assay to test immunity conferred by Pip1, we next tested engineered ePip1 by comparing this to wild-type Pip1. Importantly, growth of fluorescent *P. infestans* was further reduced in tissues expressing ePip1 when compared to wild-type Pip1 (**Figure 1D**), demonstrating that EpiC2B-insensitive ePip1 increases immunity against *P. infestans*.

Thus, we engineered the Pip1 immune protease from tomato to render it insensitive to protease inhibitor EpiC2B from *P. infestans* by mutating two residues. These two residues are invariant in Pip1 in both tomato (46 genomes, Zhou et al., 2022, Li et al., 2023) and potato (104 genomes, Tang et al., 2022), and these substitutions can be introduced into tomato cultivars by genome editing under the Precision Breeding Act that recently passed into law in the UK. In the absence of known Pip1 substrates, we can not exclude that some of this immunity might be caused by altered substrate specificity, but this is unlikely given the location of these two residues outside the substrate binding groove (**Figure 1A**). Durability of this immunity can be increased by introducing additional substitutions that suppress EpiC2B inhibition, or by building a multigene array that encodes different ePip1 variants. Similar structure-guided mutagenesis can be performed on Pip1 to increase immunity against other pathogens, and on other secreted immune proteases to increase immunity in other crops, such as maize and citrus. This approach enhances the natural extracellular immunity in crops that can be achieved using genome editing and offers a distinct alternative to engineering pathogen perception mechanisms.

## Acknowledgements

We thank Ursula Pyzio for excellent plant care, Sarah Rodgers and Caroline O’Brian for technical assistance. We thank Yasin Tumtas and Tolga Bozkurt for sharing the *P. infestans* 88069 tD strain and for advice on handling *P. infestans*. This work was financially supported by Biotechnology and Biological Sciences Research Council (BBSRC) 18RM1 project ‘Pip1S’ BB/S003193/1 and ERC-2020-AdG award 101019324 ‘ExtraImmune’.

## Author contributions

MS designed and performed most of the experiments with help of LA and SE. LA generated most Pip1 mutant strains. JK generated pJK157 and pJK489. TK and MK synthetized TK011. MS and RvdH wrote the article, with input from all authors.

## Supporting information

Supplementary Figures S1, S2

Supplemental Materials & Methods

