## Supplemental methods for "Enhanced late blight resistance by engineering an EpiC2B-insensitive immune protease"

***Protein complex prediction and visualization*** - The amino acid sequences of the mature proteases (VPN as first amino acids) and of EpiC2B were used as input for protein complex prediction. Protein complexes were modelled using AlphaFold-Multimer (Evans et al., 2022) via the ColabFold platform (Mirdita et al., 2022) and using the AlphaFold2_mmseqs2 notebook and default settings. Resulting models where annotated using PyMOL (The PyMOL Molecular Graphics System, Version 2.0 Schrödinger, LLC.).

***Molecular cloning*** – The Pip1 mutant carrying the C153A/C154A substitutions was ordered as a synthesized uncloned gene fragment (**Table S2,** ThermoFisher) and cloned into pICH41264 using BpiI, resulting in pJK152 (**Table S1**). Next, the Pip1* fragment from pJK152 was cloned into binary vector pJK268c (binary vector carrying p19, Paulus et al., 2020) together with fragments from pICH51288 (2x35S promoter, Engler et al., 2014), pJK002 (NtPR1a signal peptide and PIV2 intron, Grosse-Holz et al., 2018), and pICH41414 (35S terminator, Engler et al., 2014), using BsaI, resulting in pJK489 (**Table S1**), which carries the 35S:Pip1* on the T-DNA. Binary vectors containing the different Pip1 alleles where generated by modifying pJK488 (pL2M-P19-2x35S::Pip1, Kourelis et al., 2020) via "oligo stitching" (De Saeger et al., 2022). All used and generated vectors are listed in **Table S1**. Vector backbones were generated by PCR amplification of pJK488 using the Q5 High Fidelity PCR kit (New England Biolabs), following manufacturer's protocols. The amplified products were purified using the Quick PCR purification Kit (K310001 PureLink, Invitrogen). Single stranded, 60 bp DNA oligos (Sigma-Aldrich) where used as inserts. **Table S2** contains a complete list of PCR primers and oligos used as inserts. Plasmids were then build by Gibson Assembly (Gibson et al., 2010). Plasmids were transformed into home-made *E. coli* DH10B chemically competent cells through the heat-shock method (Sambrook and Russell, 2001). For the analysis of clones, single *E. coli* colonies were picked and grown overnight in 3 mL of LB medium (10g/L Tryptone, 10g/L NaCl, 5g/L Yeast extract, pH 7) with 50 µg/ml kanamycin at 37ºC. Plasmids were extracted and sent for Sanger sequencing using appropriate primers (**Table S2**). Correct plasmids were transformed into *Agrobacterium tumefaciens* AGL-1 (for high protein expression) or GV3101 (for pathogen assays) chemically competent cells through freeze-thaw (Sambrook and Russell, 2001). Transformed cells were grown overnight night at 28ºC in LB agar media containing 50 µg/ml kanamycin, 25 µg/ml rifampicin and either 50 µg/ml carbenicillin for AGL-1 cells or 50 µg/ml gentamycin for GV3101 cells. Positive colonies were confirmed through colony PCR and Sanger sequencing. A construct for expressing His-EpiC2B in *E. coli* was generated by cloning fragments from pJP001 (T7 promotor, Kourelis et al., 2020), pJK120 (OmpA-6His-TEV, Kourelis et al., 2020), pJK017 (EpiC2BΔSP, Grosse-Holz et al., 2018) and pJP002 (T7 terminator, Kourelis et al., 2020) into pJK082 (pET28b-GG, Kourelis et al., 2020), using BsaI, resulting in pJK157 (**Table S1**).

***Plant growth conditions*** – Wild-type *Nicotiana benthamiana* (LAB) seeds were sown into a 3:1 mix of soil (Sinclair Modular Seed Peat reduced propagation mix) with vermiculite (Sinclair brand Pro Medium) in 7x7 cm square pots and grown at high humidity under transparent plastic covers for 5 days. Seedlings were uncovered and grown for three weeks in the greenhouse at 80-120 µmol/m$^2$/s light and 21°C (night) and 22-23°C (day) in a 16-hour light regime. Plants were then agroinfiltrated and grown in a growth chamber at 100 µmols/m$^2$/sec light and 21°C and 50-60% relative humidity in a 16 h light regime. Plants were watered three times per week such that no pots were standing in water overnight.

***Production and extraction of Pip1 mutants from N. benthamiana*** – Mutant Pip1 proteins were produced in *N. benthamiana* via agroinfiltration. AGL-1 agrobacteria liquid cultures carrying the Pip1-mutant plasmids were grown at 28°C and 180 rpm for at least 24 hours in LB medium with the required antibiotics (50 µg/ml gentamicin or 25 µg/ml rifampicin and 50 µg/ml carbenicillin for AGL-1 cells or 50 µg/ml gentamycin for GV3101). Cells were harvested by centrifugation for 10 minutes at 3000 rpm and room temperature. The pellet was resuspended in infiltration buffer (10 mM MES, 10 mM $MgCl_2$, 100 µM acetosyringone at pH 5.7) and adjusted to an OD600 of 0.5. The cultures were then incubated for one hour. Three to four weeks old plants were used for infiltration. On each plant, the three youngest fully developed leaves were infiltrated with the agrobacteria using a 1ml syringe without the needle. Pip1 mutant proteins were extracted within apoplastic fluids from the infiltrated plants. For apoplastic fluid extraction, infiltrated leaves were harvested at 4 dpi. The harvested leaves were vacuum- infiltrated with cooled water. The fluids were collected in a falcon by centrifuging the leaves at 500 g and 4°C for 20 minutes.

***Production and purification of His-EpiC2B from E. coli*** - The His-EpiC2B expression construct (pJK157, **Table S2**) was transformed into *E. coli* strain Rossetta2(DE3) pLysS (Novagen/Merck). An overnight culture in LB medium, was diluted to a final OD600 of 0.012 in 200 - 600 ml TB medium (24 g/L yeast extract, 12 g/L bacto-tryptone, 0.5% glycerol, after autoclaving, 0.2 M/L potassium phosphate buffer [26.8 g $KH_2PO_4$ and 173.2 g $K_2HPO_4$ pH 7.6]) containing 50 µg/ml kanamycin. The cells were grown at 37°C and 220 rpm to reach an OD600 of 0.6. Gene expression was induced with 1 mM isopropyl β-D-1-thiogalactopyranoside (IPTG) and further incubation overnight. To collect the secreted protein, the culture was centrifuged at 10,000 × *g* at room temperature for 30 minutes. The supernatant was filtered using a vacuum system with 0.22 µm pore size (Merck Millipore, Stericup Quick Release-GP). The filtered supernatant was then incubated for at least 3 hours with 1 mL nickel-charged resin (Qiagen, Ni-NTA Superflow) per 50 ml supernatant at 4°C on a rotator. The resin was then transferred to a purification column and washed with 10 volumes of purification buffer (50 mM Tris-HCl pH 7.5, 150 mM NaCl, 10 mM imidazole), followed by 10 volumes of washing buffer (50 mM Tris-HCl, 150 mM NaCl, 50 mM imidazole, pH 8). Finally, the protein was eluted with 5 volumes of elution buffer (50 mM Tris-HCl, 150 mM NaCl, 250 mM imidazole, pH 8). The eluate was concentrated using a centrifugal filter with 3 kDa cut-off (Merck Millipore, Amicon Ultra-15 centrifugal filter units) and a buffer exchange to 50 mM sodium acetate (pH 5.0) was carried out at the same time. The protein concentration was determined using the DC Protein Assay Kit from Bio-Rad for a microplate assay with a BSA standard following the manufacturers protocol. Protein purity was verified by running the protein on 15% SDS-PAGE gels followed by Coomassie staining.

***Synthesis of TK011*** - Commercially available E64 Azide (Toronto Research Chemicals, 1 mg, 2.7 µmol, 1 eq.) was dissolved in acetonitrile (100 µL). A portion of an aqueous 100 mM CuI solution (16.2 µL, 1.6 µmol, 0.6 eq.) and DIPEA (1.9 µL, 10.8 µmol, 4 eq.) were added. Commercially available BDP-FL-alkyne (Lumiprobe, 5 mg) was dissolved in 100 µL of acetonitrile and a portion (35.7 µL, 5.4 µmol, 2 eq.) was added to the reaction solution. The solution was stirred at room temperature for 16h after which reaction control by LC-MS indicated the completion of the reaction. The desired product was isolated from the solution by injection into a preparative HPLC equipped with a RP-C18 column and run at a flow of 20 mL/min and the following gradient program (all solvents contained 0.1% (v/v) TFA): 90% $H_2O$/10% ACN to 45% $H_2O$/55% ACN in 2 minutes, to 42% $H_2O$/58% ACN in 25 minutes. Product-containing fractions were pooled and lyophilized to yield 1.6 mg (84%) of TK011. LC-MS (ESI): tR = 8.51 min, m/z calculated for $C_{33}H_{46}BF_2N_8O_6$ $[M^+H]^+$ 699.36, found 699.14.

***Activity labelling and inhibition assays*** - For inhibition assays, the extracted apoplastic fluids containing Pip1 or Pip1 mutant proteases were preincubated with purified EpiC2B or E64 (100 mM) or DMSO on reaction buffer (50 mM NaAc pH5, 10 mM DTT) for 30 minutes at room temperature on an orbital shaker. After the preincubation, the fluorescent probe TK011 (2 µM) was added to the reaction and incubated for further 3 hours and protected from light exposition. To precipitate the proteins, 4 volumes of cold acetone was added to each reaction. After vortexing, the samples were centrifuged for 5 minutes at room temperature and 13.000 x*g*. The supernatant was removed, and the pellet was air dried. Proteins were resuspended in 15 µl MilliQ water and then 15 µl 4 x gel loading buffer was added. The samples were boiled for 5 minutes and 7 µl were loaded on a 15% SDS gel. The gel was run at 180 V for about 2 hours covered from light exposition. The fluorescence signal of the gels was scanned in a Typhoon FLA 9000 (GE heathcare) with default Cy2 settings and automatic PMT. The gels were then used either for western blotting (see below) or stained with Coomassie (InstantBlue Coomassie protein stain, Abcam) as a loading control. The fluorescence signal was quantified using the Fiji platform (Schindelin et al., 2012).

***Western blot analysis*** – Proteins were transferred onto a polyvinylidene difluoride (PVDF) membrane using the TransBlot Turbo system (Bio-rad, Hercules, CA). The membrane was blocked overnight at 4°C in 5% w/v skimmed milk in tris-buffered saline (TBS) with 0.01% v/v Tween-20 (TBS-T). Pip1 mutants were detected using an α-Pip1 antibody (Tian et al., 2007). The membrane was incubated for 1 h with 1/1000 α-Pip1 antibody in 2.5% w/v skimmed milk in TBS-T. After washing the membrane with TBS-T, it was incubated for 1h with α-rabbit-HRP (Invitrogen). Chemiluminescent signals were detected using the SuperSignal™ West Femto Maximum Sensitivity Substrate (Thermo Fisher Scientific, Waltham, MA, USA).

***Phytophthora infestans* infection assays –** Infection assays were conducted on *N. benthamiana* leaves agroinfiltrated with GV3101 strains. *P. infestans* strain 88069td (Chaparro-Garcia et al., 2011) was grown in Rye Sucrose-V8 Agar (RSA-V8) (Huang et al., 2020) plates in the dark for 12 days at 18°C. Sporangia were harvested from RSA-V8 plates by adding cold water to the plates and zoospores were collected after 4 hours of incubation at 4°C. Three-week-old *N. benthamiana* leaves were agroinfiltrated side by side with two different strains. At 3dpi, the leaves were detached and placed on wet paper towels. The leaves where drop-inoculated with 10 µL droplets containing solution of 500-800 zoospores/ml. The droplets were applied onto abaxial sides of the detached leaves. One droplet/leaf half was placed on total 12 leaves per box were inoculated. Boxes where sealed and incubated in the dark at 18°C. Infected leaves were monitored between 8 and 11 days-post-inoculation and growth was documented once the hyphae had grown beyond the inoculation droplet but within the agroinfiltrated zone. Hyphal growth was documented using a Typhoon FLA 9000 scanner (GE heathcare) with default Cy3 settings and 300 PMT in all cases. Only leaves with evident hyphal growth in both sides of the leaf were used for quantification. The area covered by the hypha was measured using the red fluorescent signal and with Fiji platform (Schindelin et al., 2012).

**Table S1** Plasmids used in this study

| Plasmid | Description | Reference |
|---|---|---|
| pJK050 | pL2M-2x35S::P19 | (Kourelis et al., 2020) |
| pJK488 | pL2M-P19-2x35S::Pip1 | (Kourelis et al., 2020) |
| pJK489 | pL2M-P19//2x35S::NtPR1a-Pip1ΔSP^C153A,C154A | This work |
| pJK152 | pL0M-nsCDS1-Pip1ΔSP^C153A,C154A | This work |
| pJK157 | pET28b-T7::OmpA-6xHis-TEV-EPIC2BΔSP | This work |
| pLA2 | pL2M-P19-2x35S::Pip1_D147R_V150E | This work |
| pLA4 | pL2M-P19-2x35S::Pip1_D267E_H270E | This work |
| pLA7 | pL2M-P19-2x35S::Pip1_R284E_N286R | This work |
| pLA8 | pL2M-P19-2x35S::Pip1_D314R | This work |
| pLA9 | pL2M-P19-2x35S::Pip1_L338D_V339Y | This work |

**Table S2** Primers and oligos used in this study

| Oligo | Sequence (5'-3') | Description |
|---|---|---|
| oMS1 | CGCGGCTATAGAAGGAGCAT | Sequencing primer |
| oMS2 | GGAGAGGACCATTTGGAGAGGACACGT | Sequencing primer |
| oMS3 | GATTAGCATGTCACTATGTGTGCATCC | Sequencing primer |
| oMS4 | AGTGTGACTGAAGTGCCGAA | Sequencing primer |
| oMS6 | TATGCGTCGTTCCCTACTGC | pLA9 backbone generation |
| oMS7 | GAGAGGGAGTGTCACAGGAGTCAAGCGTCAAGGT GAATGTGGATGTTGTTGGGCATTTTC | Insert of pLA2 |
| oMS9 | ATTTCTGTTGGTATTGCGGCTAACGAAGAGTTTGAA ATGTACGGAAGTGGAATATATGAT | Insert of pLA4 |
| oMS10 | GAATATATGATGGAAGTTGTAATTCTGAACTGCGTC ATGCAGTGACAGTTATAGGTTATG | Insert of pLA7 |
| oMS11 | ATTGGATAGTGAAGAATTCATGGGGGAGTCGATGG GGTGAGGAAGGATATATGAGAATTG | Insert of pLA8 |
| oMS12 | GTTGATGGTGGCCATTGTGGCATTGCAGATTATGC GTCGTTCCCTACTGCTTAAGCTTCT | Insert of pLA9 |
| oMS13 | CTCCTGTGACACTCCCTCTC | pLS2 backbone generation |
| oMS14 | GGATGTTGTTGGGCATTTTC | pLA2 backbone generation |
| oMS17 | GCCGCAATACCAACAGAAAT | pLA4 backbone generation |
| oMS18 | ACGGAAGTGGAATATATGAT | pLA4 backbone generation |
| oMS20 | ACAACTTCCATCATATATTC | pLA7 backbone generation |
| oMS23 | AGTGACAGTTATAGGTTATG | pLA7 backbone generation |
| oMS24 | TGAATTCTTCACTATCCAAT | pLA8 backbone generation |
| oMS25 | GGAAGGATATATGAGAATTG | pLA8 backbone generation |
| oMS26 | CCACAATGGCCACCATCAAC | pLA9 backbone generation |
| Pip1* | TTGAAGACAAAGGTCGCAACTTAAAAGAATTATCCA TGCTTGAAAGGCATGAAAATTGGATGGTTCATCATG GACGTGTATACAAAGATGATATAGAAAAAGAACACC | ORF encoding Pip1 with C153A/C154A substitutions |

| | |
|---|---|
| | GCTTCAAGACATTCAAGGAAAACGTTGAGTTCATCG<br>AATCTTTCAACAAGAACGGAACTCAACGTTATAAGC<br>TAGCCGTCAATAAATATGCTGATCTGACCACTGAG<br>GAATTCACAACATCATTTATGGGGCTCGACACTTCA<br>TTACTATCCCAGCAAGAATCAACAGCTACAACAACG<br>TCCTTTAAATATGATAGTGTGACTGAAGTGCCGAAT<br>AGCATGGACTGGAGAAAGAGAGGGAGTGTCACAG<br>GAGTCAAGGATCAAGGTGTATGTGGAGCTGCATGG<br>GCATTTTCTGCGGCCGCGGCTATAGAAGGAGCATA<br>TCAAATTGCAAACAACGAACTAATCTCTCTTTCAGA<br>GCAACAACTATTGGATTGCTCTACGCAGAACAAAG<br>GCTGCGAGGGCGGACTAATGACTGTCGCTTACGAC<br>TTCTTACTTCAGAACAATGGCGGGGGCATTACCAC<br>AGAGACTAACTATCCTTATGAAGAAGCTCAAAATGT<br>TTGCAAGACCGAACAACCAGCAGCAGTTACTATCA<br>ATGGCTACGAAGTCGTACCATCTGACGAGTCATCG<br>TTGTTGAAAGCTGTAGTGAATCAGCCTATTTCTGTT<br>GGTATTGCGGCTAACGATGAGTTTCATATGTACGG<br>AAGTGGAATATATGATGGAAGTTGTAATTCTAGACT<br>GAATCATGCAGTGACAGTTATAGGTTATGGGACAA<br>GTGAAGAAGATGGTACAAAATATTGGATAGTGAAG<br>AATTCATGGGGGAGTGATTGGGGTGAGGAAGGATA<br>TATGAGAATTGCTAGAGATGTTGGAGTTGATGGTG<br>GCCATTGTGGCATTGCAAAGGTTGCGTCGTTCCCT<br>ACTGCTTGAGCTTTTGTCTTCAA |
