## Supplemental figures for "Enhanced late blight resistance by engineering an EpiC2B-insensitive immune protease"

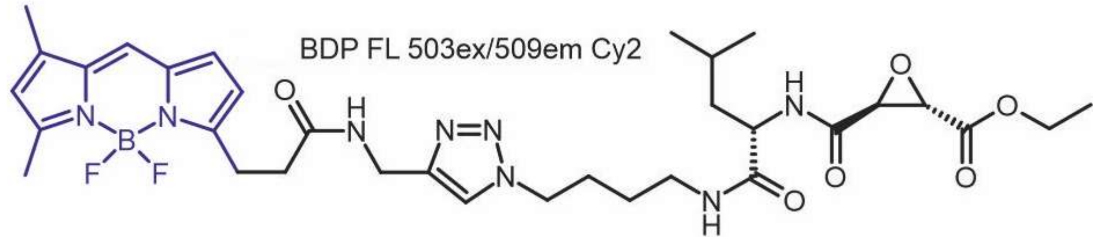


**Figure S1** TK011 is an E-64-based fluorescent probe.
Structure of the TK011 probe consisting of a E64-azide linked to an BDP-FL-alkyne (blue).

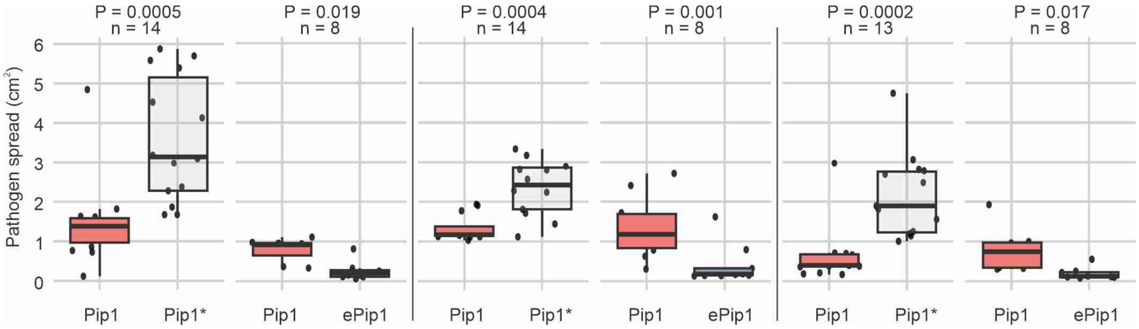


**Figure S2** Overexpression of ePip1 in *N. benthamiana* restricts *P.infestans* spread.
Three independent replicates of the experiment are shown. Data of the second replicate was used in the main figure. P values correspond to paired *t*-test for n=x biological replicates.
